# MR Spectroscopy without Water Suppression using the Gradient Impulse Response Function

**DOI:** 10.64898/2026.01.16.699878

**Authors:** James B. Bacon, Peter Jezzard, William T. Clarke

**Author notes:** Corresponding Author: William T. Clarke, FMRIB Centre, John Radcliffe Hospital, Headington, Oxford, OX3 9DU, United Kingdom.

## Abstract

**Purpose:** Non-water-suppressed proton spectroscopy, ^1^H-MRS, is desirable, as retaining the strong water resonance can facilitate automated online data corrections, internal concentration referencing, and monitoring of line narrowing effects in functional MRS. Removal of the water suppression module can also mitigate magnetization transfer effects and slightly reduce the minimum achievable TR and total RF power deposition. However, water suppression is typically considered essential due to eddy current-induced antisymmetric sidebands on the water resonance that distort the spectral baseline and obscure metabolite signals.

**Theory and Methods:** The Gradient Impulse Response Function (GIRF) was used to predict time-dependent magnetic field perturbations during the FID that generate the artefactual sidebands. The GIRF was measured in a one-time calibration, independent of spectroscopy acquisitions, enabling post-processing correction of the sidebands without sequence modification or additional dedicated hardware. GIRF-corrected non-water-suppressed single-voxel-spectroscopy (SVS) was compared to otherwise identical water-suppressed acquisitions in eight participants at 3T using semi-LASER and MEGA-PRESS sequences.

**Results:** Across participants, GIRF correction reduced sideband amplitudes to levels comparable with the spectral baseline, enabling recovery of the underlying metabolite signals. Systematic increases in quantified metabolite concentrations were observed relative to water-suppressed acquisitions, consistent with water-suppression-induced magnetization transfer effects. Total creatine exhibited the largest increase, with enhancement ratios of 1.069±0.039 for MEGA-PRESS and 1.535±0.160 for semi-LASER acquisitions.

**Conclusion:** Gradient-induced artefactual sidebands in non-water-suppressed MR spectroscopy can be effectively corrected using the GIRF to predict time-dependent magnetic field perturbations during the FID. In principle, the approach extends to other SVS sequences and field strengths following appropriate GIRF calibration.

## Introduction

Water suppression in proton spectroscopy, ^1^H-MRS, is generally considered essential, as antisymmetric sidebands on the water resonance strongly distort the spectral baseline and obscure the metabolite signals. These antisymmetric sidebands arise from time-dependent field perturbations during the FID, which are themselves caused by eddy currents and mechanical vibrations induced in the MRI hardware during gradient switching[1–3]. However, acquiring ^1^H-MRS data without water suppression offers several potential advantages.

First, removing the RF pulses required for water suppression leads to a small reduction in the total RF power deposition[4,5], shortens the minimum achievable TR, and minimizes changes in the observed amplitude of metabolite resonances through magnetization transfer effects[6–8]. Second, retaining the strong water signal may facilitate prospective frequency correction and automated retrospective phase and frequency correction, even under low SNR conditions[9], as well as correction (or identification) of motion corrupted transients in motion-sensitive diffusion techniques[10]. Third, the water signal may serve as an internal concentration reference for metabolite quantification[11], and can be used to track BOLD (line-narrowing) effects in functional MRS[12]. Finally, downfield labile peaks may remain detectable without water suppression[13]. Accordingly, several previous studies have sought to compensate for these artefactual sidebands without using water suppression[14].

Nixon *et al*.[1] proposed a theoretical model in which the spectral sidebands arise from exponentially damped sinusoidal phase modulations of the NMR signal. These modulations originate from Lorentz force-induced deformations of the gradient coil windings, leading to oscillations, thought to be mechanical[2], that perturb the static B_0_ field. Such frequency-dependent modulations are not addressed by vendor-implemented pre-emphasis schemes, which model the eddy currents solely as sums of exponential decays[15,16]. Using a model comprising 40-50 frequency-dependent terms, Nixon *et al*. demonstrated compensation of the artefactual sidebands at isocenter through the application of a predetermined correction waveform stored in local memory. However, their analysis indicated that complete suppression of sidebands away from isocenter would require damped sinusoidal pre-emphasis of the linear gradients, which could not be realized within their available hardware constraints.

Alternatively, two-scan approaches[2,5,17,18] exploit the dependence of sideband appearance on the applied gradients and the central peak state. By acquiring paired scans with opposite gradient polarity or frequency-selective inversion (excluding the central water peak, known as metabolite cycling) and subsequently summing or subtracting the pair, the sidebands can be cancelled. However, the requirement of paired acquisitions for each measurement either increases the total scan time or reduces the temporal resolution when the overall scan duration is fixed. This imposes additional acquisition constraints, particularly when combined with advanced MRS techniques such as J-difference editing, functional MRS or diffusion-weighted MRS. Furthermore, two-scan techniques increase sequence complexity and RF duty cycle and may remain sensitive to participant motion and B_0_ instability. Although adiabatic inversion pulses used in metabolite cycling substantially reduce sensitivity to B_1_^+^ inhomogeneity when the adiabatic condition is satisfied, inversion efficiency remains frequency dependent and may require correction under certain conditions[9].

Further alternatives to non-water-suppressed spectroscopy include sequence optimization and post-processing strategies aimed at reducing sidebands[4,19]. However, these approaches are typically sequence- and hardware-specific and generally achieve a partial reduction rather than complete suppression of the sidebands.

Other than the approach proposed by Nixon *et al*.[1], which was limited by hardware constraints, no previous method has explicitly sought to characterize and compensate for the system imperfections responsible for the gradient-induced sidebands in non-water-suppressed spectroscopy. Consequently, existing approaches are not readily applicable to MRS sequences used in routine practice without substantial modification. As a result, water suppression[20–22] remains the common practice, with Klose’s method[23] employed to correct sidebands on the metabolite resonances themselves.

To address these limitations and to introduce a generalized method for water-unsuppressed ^1^H-MRS, this study proposes an approach in which gradient-induced field perturbations are characterized using the Gradient Impulse Response Function (GIRF)[24]. The GIRF provides an accurate system-level model of the gradient response, capturing eddy current-related and other dynamic field effects. For a sequence with known gradient waveforms, this enables a post-processing prediction of the B_0_ perturbation during the FID and subsequent correction of the resulting artefactual sidebands. The GIRF is acquired in a one-time calibration scan, independent of spectroscopy acquisition, enabling non-water-suppressed single-voxel spectroscopy (SVS) that is generalizable across sequences, protocols and spatial locations, without the need for additional dedicated hardware.

### Theory

Introduced by Vannesjo *et al*.[24], the GIRF provides a system-level description of the gradient-induced magnetic field response in MR systems. Under the assumption of linear time-invariance (LTI), the relationship between an applied gradient waveform and the resulting magnetic field perturbation is fully characterized by the impulse response of the system.

In practice, the gradient system comprises three nominal input channels corresponding to the gradient axes 𝑗 ∈ {𝑥, 𝑦, 𝑧}. When a gradient waveform is applied along one of these axes, the generated magnetic field reflects both the prescribed waveform and the intrinsic dynamic behavior of the gradient system, including eddy currents and the associated mechanical vibrations. These effects which give rise to the artefactual sidebands observed in spectroscopy, can be well approximated as linear mechanisms[1,25,26], and are therefore incorporated into the GIRF framework.

The resulting gradient-induced field is spatially varying and includes both self-response and cross-response components. It may therefore be represented using a spherical harmonic spatial basis set indexed by 𝑚[27]. The system response is thus expressed as[28]:

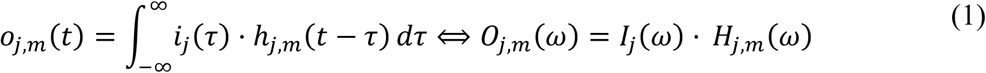

where 𝑖_j_(𝑡) denotes the input gradient waveform applied along axis 𝑗, 𝑜*_j,m_*(𝑡) represents the time-dependent coefficient of the 𝑚-th spherical harmonic field component arising from that input, and ℎ*_j_*_,*m*_(𝑡) is the corresponding impulse response. 𝐼*_j_*(𝜔), 𝑂*_j,m_*(𝜔) and 𝐻*_j,m_*(𝜔) denote their respective Fourier transforms.

For a given spherical harmonic term 𝑚, the total field may be obtained by summing 𝑜*_j,m_*(𝑡) over all gradient axes 𝑗. In the present formulation, however, axis-specific contributions are retained until evaluation at the voxel center, since the quantity of interest is the total accumulated phase error at that location.

Once measured, the GIRF can be used to predict the gradient-induced magnetic field evolution throughout an MR pulse sequence, including during the readout[29,30], in response to a known gradient input. This enables prediction of the frequency-dependent field perturbations during the FID, as illustrated in Figure 1 for the self-response term corresponding to an x-directed input.

**Figure 1:**
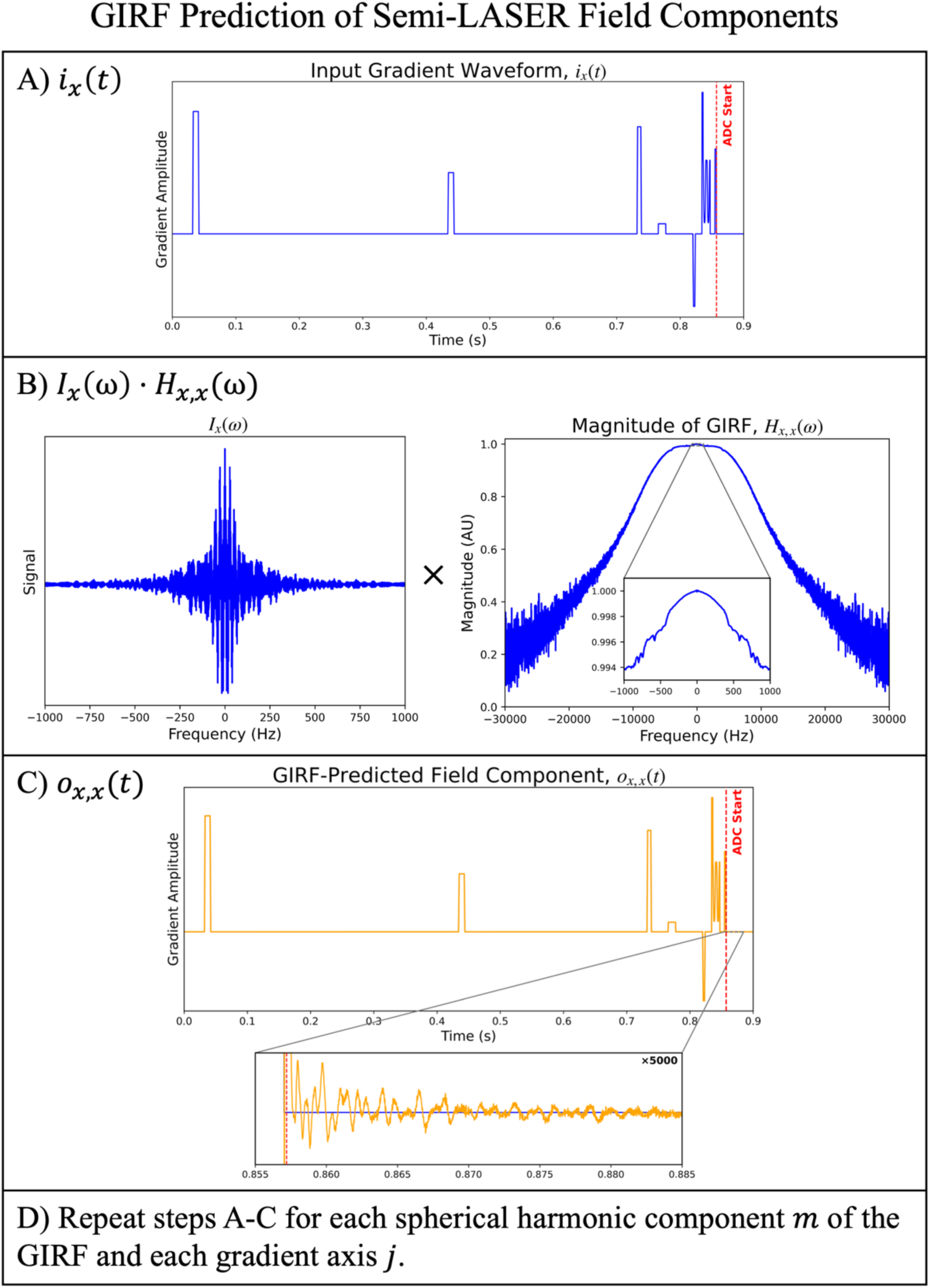
Prediction of the frequency-dependent field perturbations during the FID for a semi-LASER sequence using the GIRF. A)-C) Prediction of the x-directed self-response field component, 𝑜*_x,x_*(𝑡), by A) simulation of the input gradient waveform, 𝑖*_x_*(𝑡); B) multiplication by the self-response term of the GIRF in the frequency domain, 𝐻*_x,x_*(𝜔); and C) inverse Fourier transform to obtain 𝑜*_x,x_*(𝑡). D) To predict all field components, this procedure is repeated for each spherical harmonic component 𝑚 of the GIRF and for each gradient axis 𝑗.

In the present work, these predicted field perturbations are used to calculate the phase modulation of the NMR signal at the voxel location. The artefactual sidebands are subsequently corrected by removing this predicted phase from the measured complex signal during the readout.

## Methods

The GIRF was measured on a 3T Prisma scanner (Siemens Healthineers, Erlangen, Germany) in a one-time calibration scan using an open-source, phantom-based method[31], previously optimized for high spectral resolution[32]. An optimized thin-slice approach was employed, applying 18 positive and 18 negative triangular input gradients at four offset thin slices within the phantom[33,34]. An 11 × 11 elliptical 2D phase encoding scheme[27] was used within each slice to assess in-plane spatial variation of the GIRF, and spherical harmonic terms up to first-order were included in the subsequent sideband correction. The zeroth-order (B_0_) cross terms and self-response terms used in the correction had a spectral resolution of 5 Hz, whereas the first order-cross terms were represented at a reduced spectral resolution of 10 Hz due to the lower SNR available to these components. The full implementation of this method is available at https://github.com/jbbacon/GIRF_PE_Python.

A NiCl_2_-doped water phantom (deionized H_2_O, NiCl_2_: 5.5 mM, NaCl: 30.0 mM) was used for GIRF calibration, as it produces a single resonance peak, thereby avoiding contamination of the B_0_-cross term by additional resonances that would otherwise propagate into the corrected SVS spectra. The phantom was housed in an 11 cm diameter spherical HDPE casing (Spherical Phantom Application Research Kit (SPARK), Gold Standard Phantoms, Sheffield, UK), with the reduced diameter selected to minimize B_1_^+^ inhomogeneity. Any spherical water phantoms exhibiting a single resonance peak are in principle also suitable for GIRF calibration.

In vivo SVS measurements were performed on the same scanner in eight healthy participants (32.1±12.3 years, 71.6±14.7 kg, 5 male). Data from each participant were acquired on separate days, all distinct from the day of GIRF measurement. Data was acquired under a technical development protocol approved by local ethics and institutional committees. Measurements were also acquired in a uniform, metabolite-containing SPECTRE phantom (Gold Standard Phantoms, Sheffield UK).

A vendor-harmonized semi-LASER sequence from the CMRR spectroscopy package[35] (20 mm isotropic voxel; TE/TR: 30/5000 ms; bandwidth: 6000 Hz; GOIA-WURST pulses, 64 averages) was used in the motor cortex, and a MEGA-PRESS sequence, locally modified from the CMRR spectroscopy package[36], (25 × 20 × 25 mm^3^ voxel; TE/TR: 68/1500 ms; bandwidth: 4000 Hz, 160 averages) was used in the early-visual cortex (oblique orientation). Voxel locations and orientations have been made available for all participants in the Supporting Information (Table S1). Non-water-suppressed spectra were acquired with VAPOR[20] water suppression disabled, while water-suppressed spectra with VAPOR enabled were acquired as a comparison. Outer-volume suppression (OVS) was enabled in both cases. To assess the potential effects of system heating on the GIRF correction, the non-water-suppressed semi-LASER spectra were reacquired approximately 25 minutes after initial acquisition, and immediately following a 6-minute diffusion tensor imaging sequence (UK Biobank protocol[37]), selected to induce system heating through a high gradient duty cycle.

For both water-suppressed and non-water-suppressed acquisitions standard preprocessing steps were applied using FSL-MRS[38], version 2.4.9. These steps included RF coil combination, retrospective correction of frequency and phase drifts, and signal averaging[39]. For the water-suppressed data only, additional preprocessing included correction of eddy current effects using the Klose method[23] and removal of residual water using HLSVD. Automated phasing was performed using the NAA resonance for the water-suppressed data and using the water resonance for the non-water-suppressed data.

Eddy current-induced artefactual sidebands were removed in the non-water-suppressed data as follows:

1. The gradient waveforms for each sequence were simulated using the Siemens POET pulse sequence simulation tool. Because these waveforms depend on voxel orientation, the sequences were re-simulated for each participant using the subject-specific in vivo acquisition parameters. The resulting waveforms, 𝑖*_j_*(𝑡), were extracted and used as inputs for subsequent processing.
2. The input waveforms were multiplied with the measured GIRF, 𝐻*_j,m_*(𝜔), in the frequency domain for each gradient axis 𝑗 and spherical harmonic term 𝑚, according to Equation 1. Following inverse Fourier transformation, this yielded the predicted time-dependent magnetic field components, 𝑜*_j,m_*(𝑡), throughout the sequence, including during the readout (Figure 1).
3. The phase coefficients, 𝑘*_j,m_*(𝑡), were computed by time-integrating the predicted field components over the readout window up to each time point:

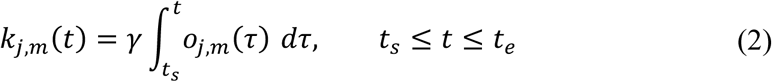

where *γ* is the proton gyromagnetic ratio, 𝑡*_s_* and 𝑡*_e_* denote the start and end of the readout period, respectively.
4. The total accumulated phase error due to gradient-induced field perturbations was then obtained as

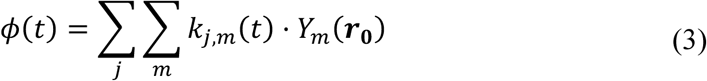

where 𝑌*_m_*(𝒓) denotes the *m*-th spherical harmonic basis function evaluated at position 𝒓, and 𝒓_𝟎_ represents the voxel-center coordinates defined in the scanner reference frame.
5. The measured FID was corrected by subtracting the predicted phase error, 𝜙(𝑡), from the phase of the measured complex signal at each time point following temporal alignment.

For the GIRF-corrected non-water-suppressed spectra (Figure 2), the water resonance was modeled separately and removed prior to metabolite quantification. This modelling was performed using FSL-MRS with a dedicated water basis set and a Voigt lineshape model. The fit was restricted to a frequency range of -10 to 4.55 ppm, rather than spanning the full water peak, to ensure accurate modelling in the region overlapping the metabolite resonances, while minimizing interference with the spectral baseline (Figure 3A). The fitted water signal was subsequently subtracted from the spectra. Examples of the resulting unfitted metabolite spectra, following water removal, are shown in Figure 3 B&C, with spectra from all participants provided in Supporting Information Figure S1.

**Figure 2:**
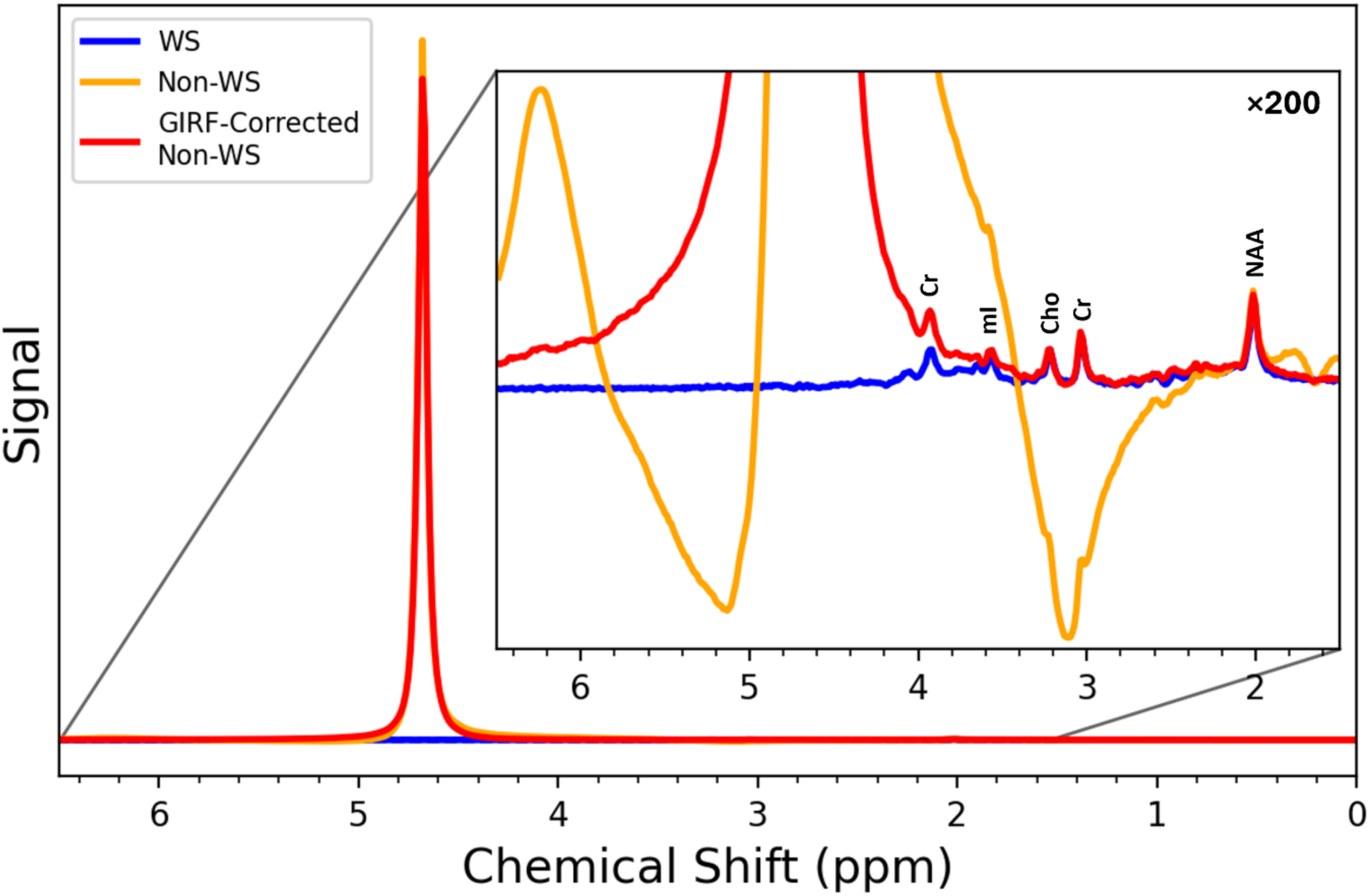
The GIRF-corrected non-water-suppressed spectrum (GIRF-Corrected Non-WS, red) shown alongside the uncorrected non-water-suppressed spectrum (Non-WS, orange) and the water-suppressed reference spectrum (WS, blue) for a representative participant acquired using the semi-LASER acquisition. The GIRF correction retains the strong water resonance while substantially reducing the artefactual antisymmetric sidebands, yielding a spectrum comparable to the water-suppressed reference. Key metabolite resonances are labelled in the magnified (×200) view. Spectra from all subjects and sequences are shown in Supporting Information Figure S1.

**Figure 3:**
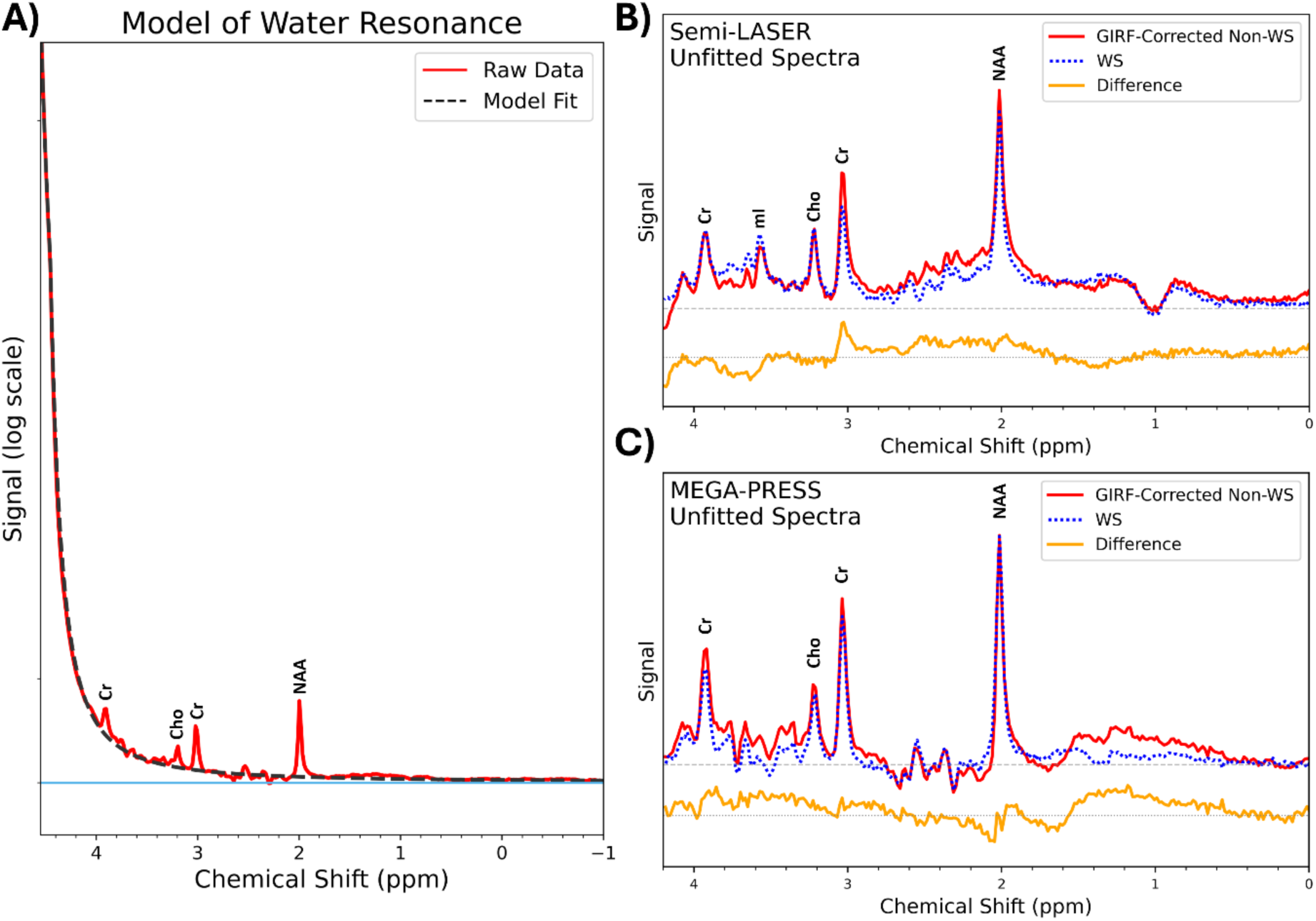
A) Modelling of the water resonance in an example MEGA-PRESS ‘off’ spectrum (editing pulses at control position[41]). The water signal is fit over a restricted frequency range without the inclusion of a baseline. B, C) Semi-LASER and MEGA-PRESS ‘off’ spectra for an example participant following subtraction of the modelled water signal from the raw data. The corresponding water-suppressed reference spectra are shown alongside the difference relative to the GIRF-corrected non-water-suppressed spectra. For visual clarity, the difference spectra are vertically offset.

Metabolite signals were fitted for both the GIRF-corrected non-water-suppressed-spectra and the water-suppressed reference spectra using sequence- and echo-time-specific metabolite basis sets simulated using FSL-MRS (Figure 4). Each basis set comprised 18 metabolites and an appropriate mobile macromolecule spectrum (semi-LASER: empirically measured using a metabolite-nulled acquisition; MEGA-PRESS: parameterized based on published literature[40]). In addition, a lipid resonance was included for the semi-LASER acquisition of one participant exhibiting substantial lipid contamination. For the MEGA-PRESS dataset, only the unedited ‘off’ spectra were fitted, as the water-resonance, sidebands, and GIRF correction are cancelled out during the editing process[41]; i.e., high quality MEGA-PRESS data can be acquired without water suppression or sideband removal . Spectral fitting was performed using a Voigt lineshape model with a “moderate” penalized spline baseline[42] over a frequency range of 0.2-4.2 ppm.

**Figure 4:**
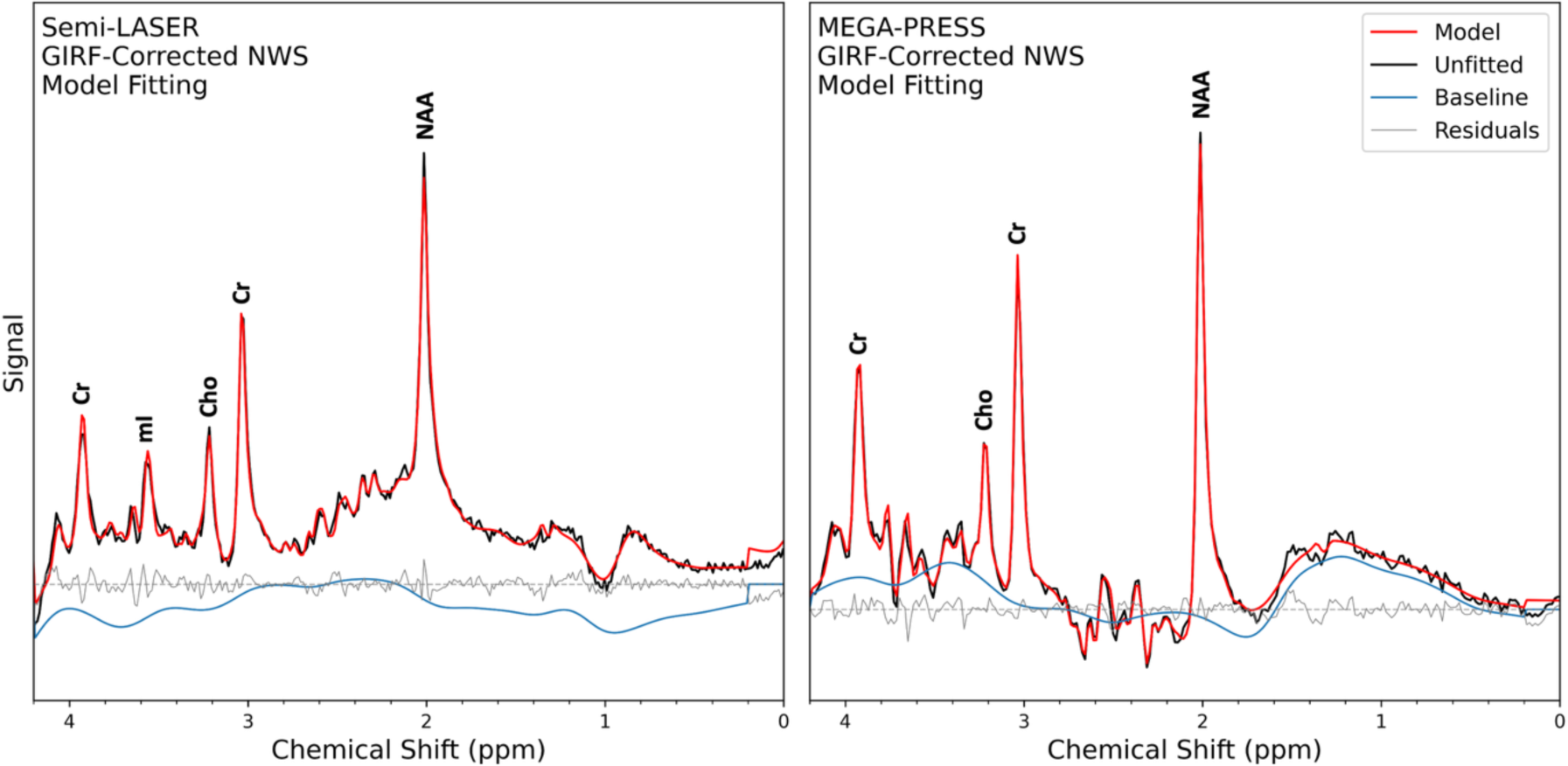
Model fitting of GIRF-corrected non-water suppressed metabolite spectra for an example participant using FSL_MRS[38], following subtraction of the modelled water signal. Spectra are shown for semi-LASER (left) and MEGA-PRESS (right) with the fitted model (red), unfitted spectra (black), fitted spectral baseline (blue) and residuals (gray).

The full pipeline including preprocessing, GIRF correction and spectral fitting is made available with example data at https://github.com/jbbacon/NWS_Spectroscopy. All processing was performed retrospectively offline in Python.

## Results

Representative examples of the fitted metabolite spectra are shown in Figure 5, alongside the corresponding water-suppressed reference spectra. Fitted spectra for all participants are provided in Supporting Information Figure S2, including the semi-LASER spectra reacquired following mild system heating (Supporting Information Figure S3). In each case, the GIRF-corrected non-water-suppressed spectra closely reproduce water-suppressed reference results, except for the creatine resonances at 3.03 and 3.92 ppm, which appear elevated in the GIRF-corrected data. Visually, agreement between the GIRF-corrected and reference-spectra is stronger for the MEGA-PRESS dataset than for the semi-LASER acquisitions.

**Figure 5:**
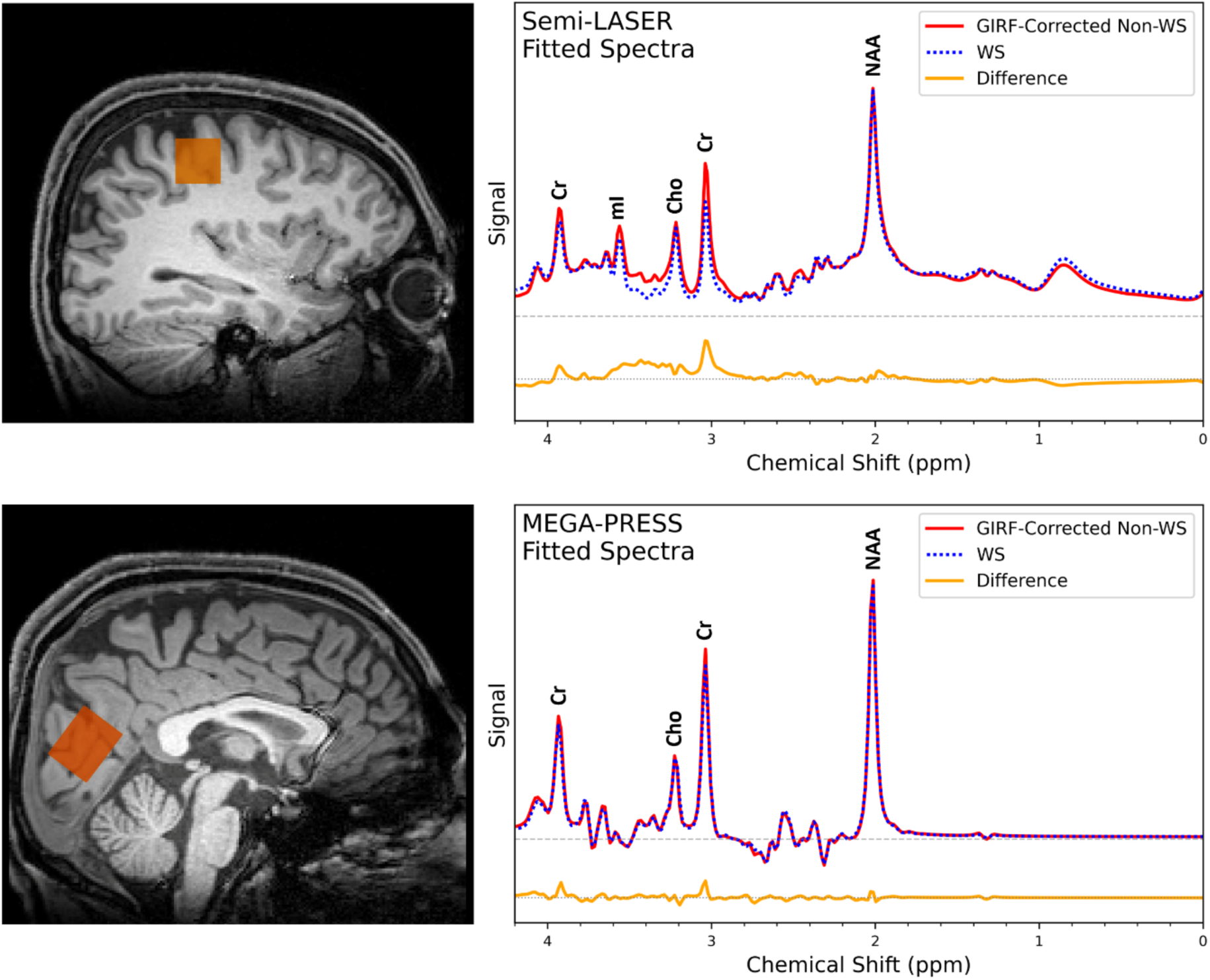
Comparison of the GIRF-corrected non-water-suppressed (red) and water-suppressed (blue) fitted spectra (baseline removed) for a representative participant. The difference spectrum (orange) is vertically offset for clarity. The corresponding voxel locations are shown on the anatomical images (left). Creatine resonances appear elevated in the GIRF-corrected spectra, with larger differences observed in the semi-LASER acquisition.

Table 1 summarizes the average concentrations of key metabolites measured across participants for each sequence acquired without water suppression. Concentrations were quantified using the retained water resonance as an internal reference and, additionally, using a separately acquired in vivo water reference scan (with RF water suppression pulses and OVS disabled), consistent with the reference approach typically used for water-suppressed acquisitions[43].

**Table 1:** Absolute concentrations (mean ± SD, n = 7) of key metabolites measured without water suppression. Quantification was performed relative to the retained water resonance (internal water reference), and to a separately acquired in vivo water reference scan (with RF water suppression and OVS disabled). Seven participants are included, as the separately acquired water reference was not obtained for one participant. Myo-inositol is not quantified for the MEGA-PRESS data due to its low signal in the ‘off’ spectrum. Abbreviations: tNAA, total NAA; tCr, total creatine; mI, myo-inositol; tCho, total choline; Glx, glutamate + glutamine.

| Metabolite | Semi-LASER<br>Concentration / mM |  | MEGA-PRESS<br>Concentration / mM |  |
| --- | --- | --- | --- | --- |
|  | Internal Water<br>Reference | Separate In Vivo<br>Water Reference | Internal Water<br>Reference | Separate In Vivo<br>Water Reference |
| tNAA | 14.88±0.97 | 13.09±0.88 | 24.67±2.93 | 13.45±1.11 |
| tCr | 10.68±1.41 | 9.41±1.31 | 14.31±1.00 | 7.82±0.30 |
| mI | 7.47±0.67 | 6.58±0.69 | - | - |
| tCho | 1.77±0.22 | 1.55±0.18 | 2.17±0.13 | 1.19 ± 0.06 |
| Glx | 7.75±1.56 | 6.84±1.45 | 15.32±2.18 | 8.35±0.83 |

Figure 6 and Table 2 summarize the ratio of the metabolite concentrations measured with and without water suppression, quantified using the same water reference. In the MEGA-PRESS data, there is strong agreement between the two approaches, except for the expected increase of total creatine which exhibits an enhancement ratio (non-suppressed to suppressed) of 1.069±0.039. The semi-LASER data shows more variability, with significant increases in total NAA, total creatine, myo-inositol, and total choline. Among these, total creatine exhibits the largest increase, with an enhancement ratio of 1.535±0.160. No significant differences were observed in the metabolite ratios for the semi-LASER data acquired before and after system heating.

**Figure 6:**
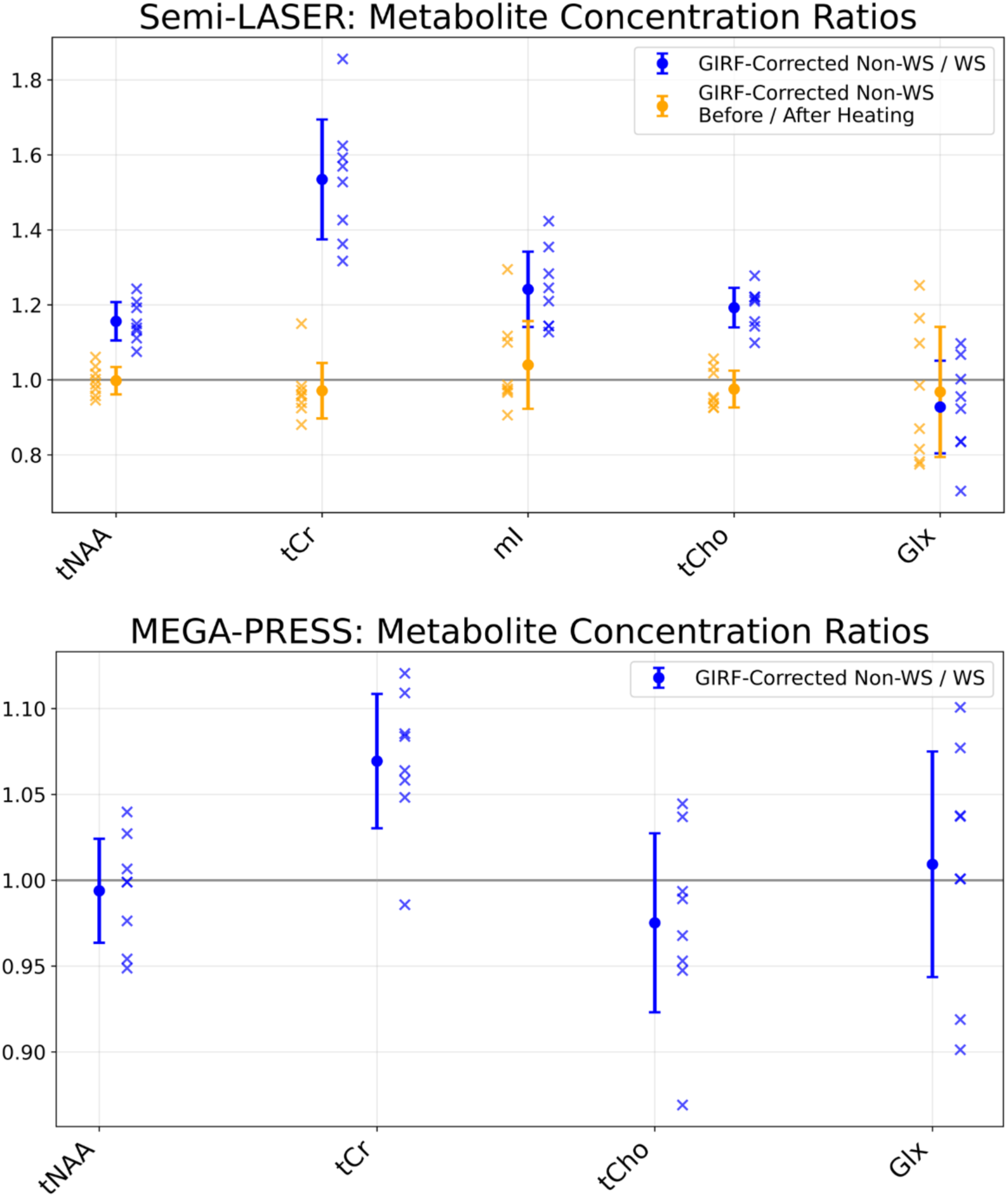
Ratios of key metabolites measured with and without water suppression (blue) and before and after system heating (orange). Top: Semi-LASER data shows strong consistency before and after system heating, but greater variability between water-suppression conditions. Bottom: MEGA-PRESS data shows strong agreement, except for total creatine.

**Table 2:** Ratios (mean ± SD, n = 8) of metabolite concentrations measured with and without water-suppression and before and after system heating. Myo-inositol is not quantified for the MEGA-PRESS data due to its low signal in the ‘off’ spectrum.

| Metabolite | Semi-LASER<br>Non-WS / WS | Semi-LASER<br>Before / After Heating | MEGA-PRESS<br>Non-WS / WS |
| --- | --- | --- | --- |
| tNAA | 1.156±0.051 | 0.998±0.037 | 0.994±0.030 |
| tCr | 1.535±0.160 | 0.971±0.074 | 1.069±0.039 |
| ml | 1.242±0.100 | 1.040±0.117 | - |
| tCho | 1.193±0.053 | 0.975±0.049 | 0.975±0.052 |
| Glx | 0.927±0.123 | 0.968±0.174 | 1.009±0.066 |

Figure 7 presents the SPECTRE phantom data, including the measured spectra following water removal, the model fits, and the final fitted metabolite spectra. The measured spectra exhibit small residual features at 2.7 ppm and 0.1 ppm in both sequence acquisitions, consistent with minor eddy currents not fully captured by the GIRF correction, resulting in slight discrepancies relative to the water suppressed reference at these frequencies. These residuals are accounted for by the spline baseline during fitting and are absent from the final fitted metabolite spectra.

**Figure 7:**
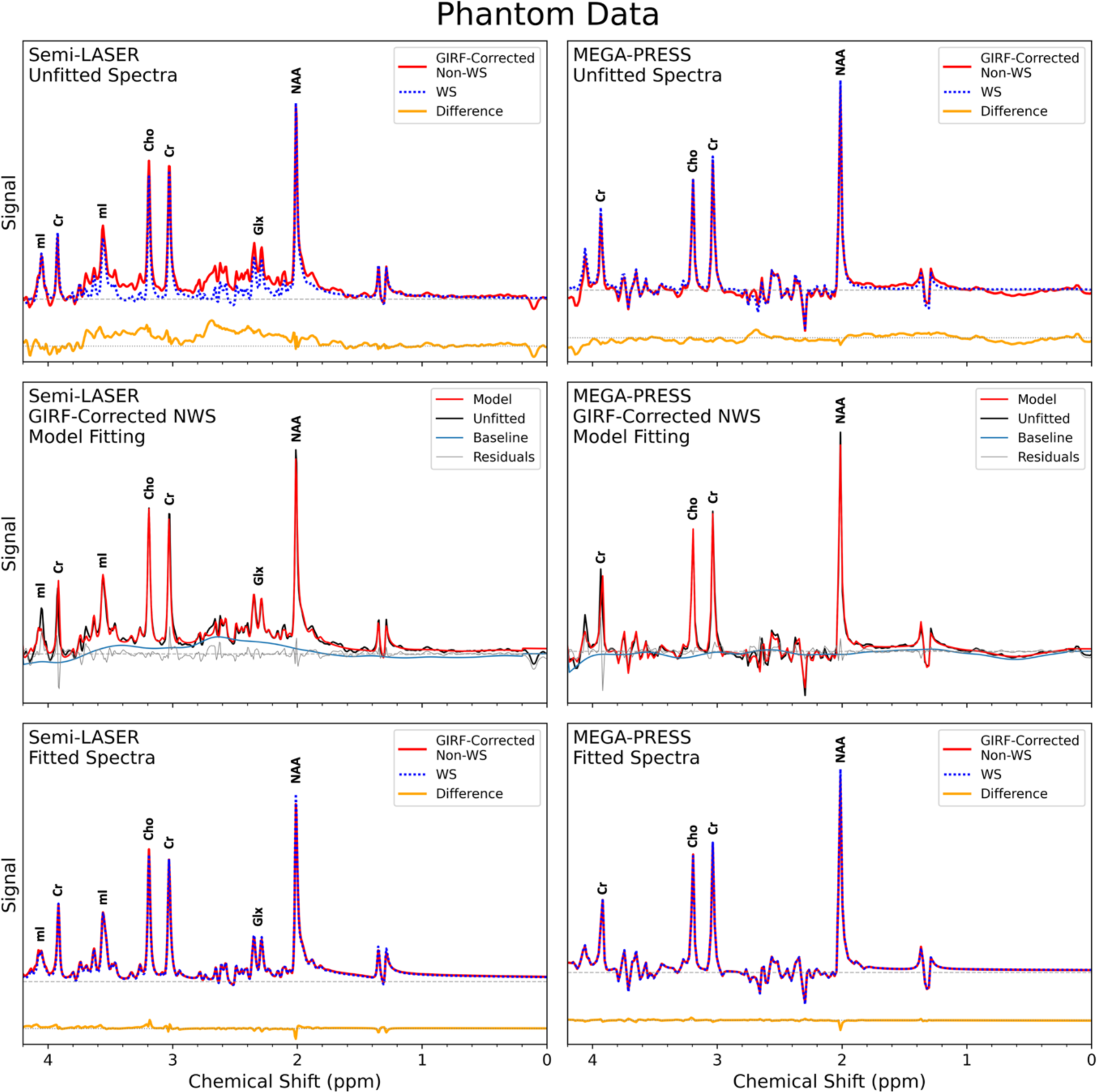
Comparison of the GIRF-corrected non-water-suppressed (red) and water-suppressed (blue) spectra acquired in the SPECTRE phantom. Top row: Measured spectra following water removal, exhibiting small residual eddy current features at 2.7 ppm and 0.1 ppm. Middle row: Model fitting of the GIRF-corrected non-water-suppressed spectra. Bottom row: Final fitted spectra, showing only minor diYerences attributable to phase misalignment associated with narrow linewidths. A 6 Hz apodization was applied to all spectra for visualization.

In contrast to the in vivo results, no visible disagreement is observed between the fitted GIRF-corrected non-water-suppressed spectra and the water-suppressed references in the phantom data. Table 3 summarizes the metabolite concentration ratios between the two approaches, with no significant differences detected other than a small increase in the total choline for the semi-LASER data.

**Table 3:** Ratios (value ± Cramér-Rao Lower Bound (CRLB) (%)) of metabolite concentrations measured with and without water suppression and before and after system heating in the SPECTRE phantom. Myo-inositol is not quantified for the MEGA-PRESS data due to its low concentration in the unedited ‘off’ spectra.

| Metabolite | Semi-LASER<br>Non-WS / WS | Semi-LASER<br>Before / After Heating | MEGA-PRESS<br>Non-WS / WS |
| --- | --- | --- | --- |
| tNAA | 0.994±3.945% | 1.004±6.252% | 0.968±3.292% |
| tCr | 1.009±4.034% | 0.999±6.489% | 0.987±3.744% |
| mI | 1.001±4.762% | 0.941±7.029% | - |
| tCho | 1.083±4.263% | 1.005±6.703% | 1.016±4.516% |
| Glx | 0.985±7.343% | 0.967±9.745% | 0.956±7.954% |

## Discussion

This study demonstrates that gradient-induced artefactual sidebands in non-water-suppressed MR spectroscopy can be substantially reduced using the GIRF to predict time-dependent B_0_ variations during the FID. Across all participants, the application of the GIRF correction reduced the amplitude of the sidebands to levels comparable to the spectral baseline, enabling recovery of the underlying metabolite signals in non-water-suppressed acquisitions.

Small residual features consistent with eddy currents were observed in some GIRF-corrected spectra (Figures 7 and S1), most prominently at 2.7 ppm and 0.1 ppm in the phantom acquisitions. Although these residuals were largely accounted for during spectral fitting using a spline baseline and did not significantly affect metabolite quantification, they highlight practical limitations of GIRF-based prediction of field perturbations.

A key limitation of this approach is that the GIRF is measured in a separate, one-time calibration, such that the correction performance depends on the scanner state at the time of the spectroscopy acquisition. This state may vary due to gradient heating or system drift between sessions. To assess sensitivity to such effects, non-water-suppressed semi-LASER spectra were reacquired following 6 minutes of high gradient duty cycle diffusion imaging, selected to induce system heating. Although this assessment was not exhaustive, metabolite concentrations remained consistent before and after heating, indicating robustness of the correction to changes in the system state within a single session. This consistency also demonstrates the repeatability of the proposed approach when applied to the same participant.

A second limitation is the trade-off between spectral resolution and SNR in the GIRF measurement. In this study, the B_0_-cross terms and self-response terms of the GIRF, which account for the majority of measured eddy current effects[1], were characterized with a spectral resolution of 5 Hz. This represents an improvement over the spectral resolution used in imaging studies[24,29,33] (≈20 Hz) and is necessary to accurately capture low-frequency field perturbations (<500 Hz) relevant to spectroscopy at 3T. However, increasing the spectral resolution by extending the temporal window used to calculate the GIRF reduces measurement SNR, particularly at higher frequencies. As a result, correction of eddy current components in the kilohertz range remained incomplete. Furthermore, spherical harmonic terms higher than first order were excluded from the correction, as these components exhibit substantially lower SNR and would increase noise propagation into the corrected spectra. For the present measurement approach, further improvements in spectral resolution are unlikely, without introducing excessive noise into the frequency band relevant for spectroscopy.

The procedure used to model and remove the water resonance in the GIRF-corrected data has not been exhaustively refined. The Voigt lineshape model employed may be insufficient to accurately characterize the water resonance at amplitudes approaching those of the metabolite signals. Imperfect water modeling could therefore introduce subtle baseline distortions, such as those observed between 0 and 2 ppm in Figure 3C and, to a lesser extent, in other subjects (Supporting Information Figure S1B).

Over longer timescales, degradation in gradient performance may affect correction accuracy, and recalibration would be required following hardware modifications, such as gradient coil replacement. Previous work has demonstrated GIRF stability within 0.1% below 10 kHz over a period of three years[29]; however the optimal recalibration interval for the present application remains unclear.

Although the small residual eddy current features were largely accounted for during spectral fitting, they may potentially be further mitigated at the acquisition stage through real-time field monitoring using a dynamic field camera. Direct measurement of the output field would replace model-based prediction and may therefore be less sensitive to inter-session variability caused by heating and system drift. In addition, field cameras enable monitoring of higher-order spherical harmonic components, typically up to third order, which could contribute to the small residuals observed in this study. Furthermore, because this approach does not rely on the LTI assumption, it may also capture non-linear behavior characteristics such as that of gradient amplifiers[30,44]. However, such hardware remains costly and is not widely available in either research or clinical settings.

When the retained water resonance was used as a concentration reference, absolute concentrations were higher than those obtained using a separately acquired in vivo water reference scan, with RF water suppression pulses and OVS disabled. This behavior is expected, as the OVS saturation bands applied in the non-water-suppressed acquisition partially attenuate the water signal near voxel boundaries and induce magnetization transfer effect within the voxel. The combined effect of OVS is therefore a reduction in the water reference signal, thereby increasing the metabolite to water signal ratio from which concentrations are derived[43]. Consequently, concentrations quantified using non-water-suppressed spectroscopy may appear elevated relative to values typically reported in literature.

Figure 6 demonstrates systematic increases in several metabolite concentrations measured without water suppression relative to reference spectra acquired with VAPOR enabled. In the MEGA-PRESS data, this effect was observed only for total creatine, whereas the semi-LASER data showed larger and more widespread increases in total NAA, total creatine, myo-inositol, and total choline, with the greatest relative increase observed for total creatine. These differences were reproducible across participants and were unaffected by mild system heating, indicating that they are unlikely to arise solely from residual gradient-induced artefacts.

We attribute these systematic differences to the water-mediated saturation transfer effects associated with the use of water suppression. When the water magnetization is manipulated using schemes such as VAPOR, saturation of the water pool can be transferred to the metabolite resonances, resulting in an apparent reduction in the measured metabolite concentrations relative to non-water-suppressed acquisitions[6]. Such effects have been reported in several previous studies, including increased metabolite concentrations when off-resonance saturation pulses are applied[7,8,45], and when non-water-suppressed spectroscopy is performed using metabolite cycling[17,18]. Across these studies, total creatine has consistently been reported as the metabolite most strongly affected by saturation transfer effects.

This interpretation is further supported by the absence of corresponding concentration differences in the SPECTRE phantom measurements. Water-mediated magnetization transfer effects are expected to be strongest in biological tissue, where the presence of many small compartments and restricted molecular mobility facilitate efficient magnetization exchange[8,45,46]. In contrast, the phantom represents a single, large, homogenous compartment with high molecular mobility; conditions under which saturation transfer effects are expected to be minimal. Consistent with this expectation, no systematic differences in metabolite concentrations were observed between water-suppressed and non-water-suppressed acquisitions in the phantom data.

The larger magnitude and more widespread increases observed in the semi-LASER data compared to the MEGA-PRESS data may reflect sequence-dependent sensitivity to magnetization transfer effects and macromolecular contributions. The semi-LASER acquisitions were performed with a shorter echo time, resulting in a substantially larger macromolecular signal contribution. Quantification in the presence of a strong macromolecular background is an inherent challenge in MRS, particularly when flexible baseline models are used[47], and this challenge may be further compounded in non-water-suppressed acquisitions, as the macromolecular signals themselves have been reported to exhibit saturation transfer effects[45,48]. The combined influence of water-mediated saturation transfer effects and macromolecular modelling limitations may therefore contribute to the larger apparent changes observed in the semi-LASER acquisitions. Future work may benefit from the use of macromolecular basis sets specifically measured for non-water-suppressed acquisitions.

## Conclusion

This work demonstrates that gradient-induced artefactual sidebands in non-water-suppressed MR spectroscopy can be effectively corrected by using the Gradient Impulse Response Function (GIRF) to predict time-dependent magnetic field perturbations during the FID. The GIRF is measured in a one-time calibration, independently of the spectroscopy acquisition, enabling post-processing correction of the sidebands without sequence modification or additional dedicated scanner hardware. While demonstrated here for two single-voxel-spectroscopy (SVS) sequences at 3T, the approach is, in principle, extendable to other SVS sequences and field strengths, following appropriate GIRF calibration.

## Data and Code Availability Statement

The data and code used in this work are made available online. Code relating to the measurement and calculation of the Gradient Impulse Response Function (GIRF) is available at https://github.com/jbbacon/GIRF_PE_Python (specific git hash 382f4e1) with data and a permanent record available at https://doi.org/10.5281/zenodo.15350583. Code and data relating to preprocessing, GIRF correction and spectral fitting is available at https://github.com/jbbacon/NWS_Spectroscopy (specific git hash 85328c6) with a permanent record available at https://doi.org/10.5281/zenodo.18262880. Data is provided after initial preprocessing for one representative participant to demonstrate the GIRF correction pipeline, and for all participants prior to spectral fitting.

## Acknowledgements

This research was funded by the Wellcome grant [225924/Z/22/Z] and supported by the NIHR Oxford Health Biomedical Research Centre (NIHR203316). The views expressed are those of the authors and not necessarily those of the NIHR or the Department of Health and Social Care. The Centre for Integrative Neuroimaging was supported by core funding from the Wellcome Trust (203139/Z/16/Z and 203139/A/16/Z). JB and PJ acknowledge support from the Vivensa Foundation. PJ also acknowledges support from the NIHR Oxford Biomedical Research Centre (NIHR203311).

This research was funded in whole, or in part, by the Wellcome Trust [Grant number 225924/Z/22/Z]. For the purpose of open access, the author has applied a CC BY public copyright licence to any Author Accepted Manuscript version arising from this submission.

## Supporting Information

Supporting Information Table S1: Voxel locations, orientations and rotations for all participant and phantom acquisitions. Voxel prescription parameters are reported in the Siemens patient-based coordinate system for a head-first supine patient position. Abbreviations: L, left; R, right; A, anterior; P, posterior; H, head; F, foot; C, coronal; T, transverse.

Supporting Information Figure S1: Unfitted spectra for all participants following water removal for A) Semi-LASER acquisitions and B) MEGA-PRESS acquisitions.

Supporting Information Figure S2: Fitted spectra for all participants for A) Semi-LASER acquisitions and B) MEGA-PRESS acquisitions. The fitted spectral baselines are also shown and have been vertically offset for visual clarity.

Supporting Information Figure S3: Fitted spectra for all participants before and after system heating in the semi-LASER acquisitions. The fitted spectral baselines are also shown and have been vertically offset for visual clarity.

