## Supporting Information for "MR Spectroscopy without Water Suppression using the Gradient Impulse Response Function"

| | Semi-LASER | | MEGA-PRESS | |
|---|---|---|---|---|
| **Participant** | **Voxel Position (mm)** | **Voxel Orientation and Rotation** | **Voxel Position (mm)** | **Voxel Orientation and Rotation** |
| **001** | L: 34.7<br>P: 8.7<br>H: 47.2 | Orientation: Coronal<br>Rotation: 0° | L: 1.0<br>P: 56.5<br>F: 3.0 | Orientation: T>C 36.9°<br>Rotation: 0° |
| **002** | L: 39.8<br>A: 9.1<br>H: 24.2 | Orientation: Coronal<br>Rotation: 0° | L: 3.7<br>P: 38.5<br>F: 17.6 | Orientation: T>C 38.7°<br>Rotation: -180° |
| **003** | L: 32.7<br>P: 2.1<br>H: 54.8 | Orientation: Coronal<br>Rotation: 0° | L: 0.0<br>P: 42.2<br>H: 14.8 | Orientation: C>T 43.4°<br>Rotation: 90° |
| **004** | L: 29.7<br>P: 13.9<br>H: 34.6 | Orientation: Coronal<br>Rotation: 0° | R: 3.1<br>P: 55.9<br>H: 0.1 | Orientation: T>C36.9°<br>Rotation: 0° |
| **005** | L: 29.6<br>P: 16.7<br>H: 50.2 | Orientation: Coronal<br>Rotation: 0° | L: 2.3<br>P: 59.1<br>H: 4.6 | Orientation: T>C 36.9°<br>Rotation: 0° |
| **006** | L: 31.9<br>A: 6.7<br>H: 56.8 | Orientation: Coronal<br>Rotation: 0° | L: 2.4<br>P: 35.3<br>H: 18.6 | Orientation: C>T 36.1°<br>Rotation: 0° |
| **007** | L: 28.3<br>P: 19.6<br>H: 30.4 | Orientation: Coronal<br>Rotation: 0° | R: 1.2<br>P: 54.6<br>F: 10.9 | Orientation: T>C 34.7°<br>Rotation: 0° |
| **008** | L: 33.2<br>A: 3.2<br>H: 46.7 | Orientation: Coronal<br>Rotation: 0° | L: 2.0<br>P: 36.6<br>H: 14.5 | Orientation: C>T 39.5°<br>Rotation: 90° |
| **Phantom** | L: 29.7<br>P: 13.9<br>H: 34.6 | Orientation: Coronal<br>Rotation: 0° | R: 3.1<br>P: 40.9<br>H: 0.1 | Orientation: T>C 36.9°<br>Rotation: 0° |

*Supporting Information Table S1: Voxel locations, orientations and rotations for all participant and phantom acquisitions. Voxel prescription parameters are reported in the Siemens patient-based coordinate system for a head-first supine patient position. Abbreviations: L, left; R, right; A, anterior; P, posterior; H, head; F, foot; C, coronal; T, transverse.*

**A)**

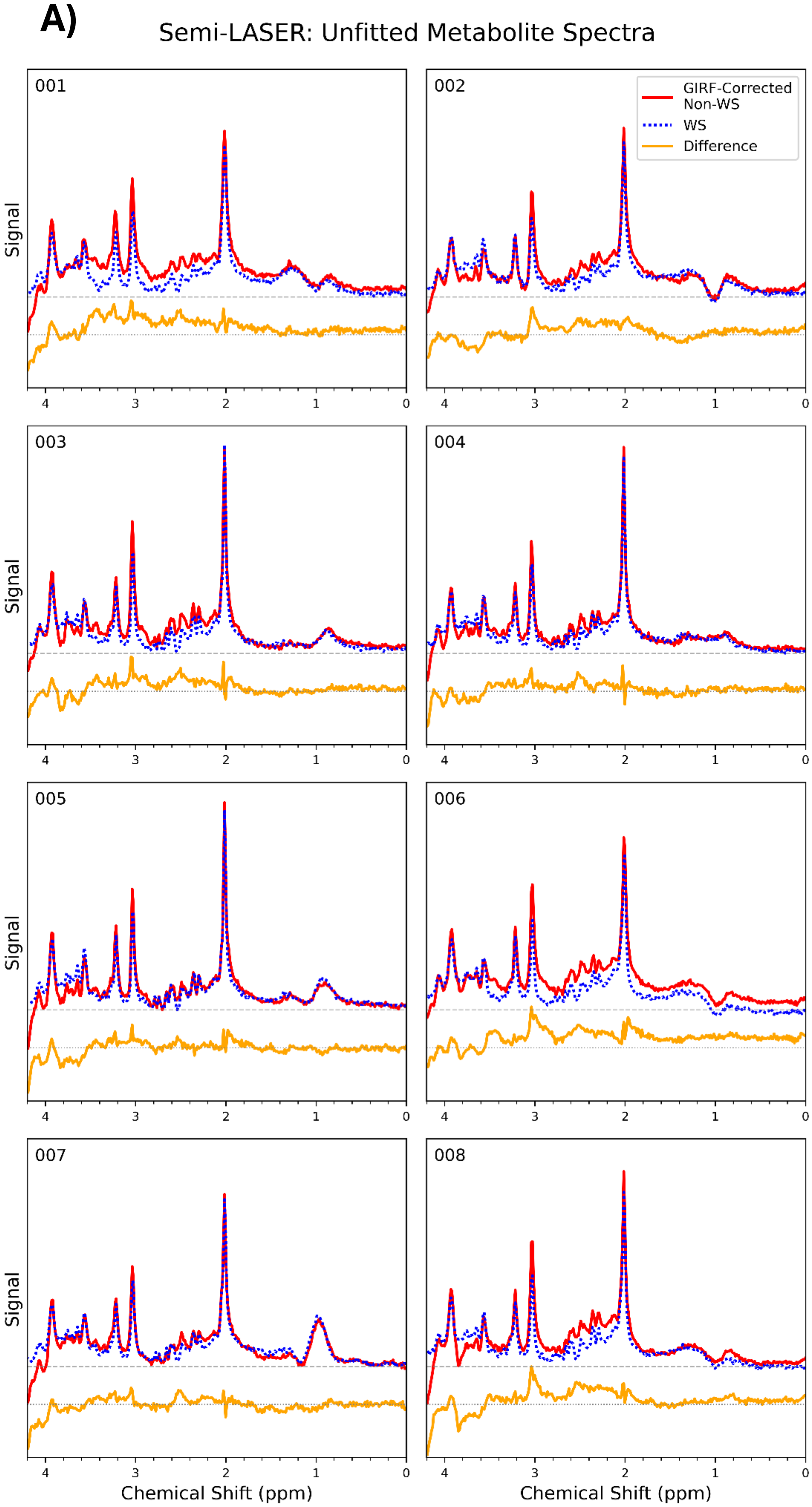

Semi-LASER: Unfitted Metabolite Spectra
001
002
003
004
005
006
007
008
GIRF-Corrected Non-WS
WS
Difference
Signal
Chemical Shift (ppm)
4
3
2
1
0

**B)** MEGA-PRESS: Unfitted Metabolite Spectra

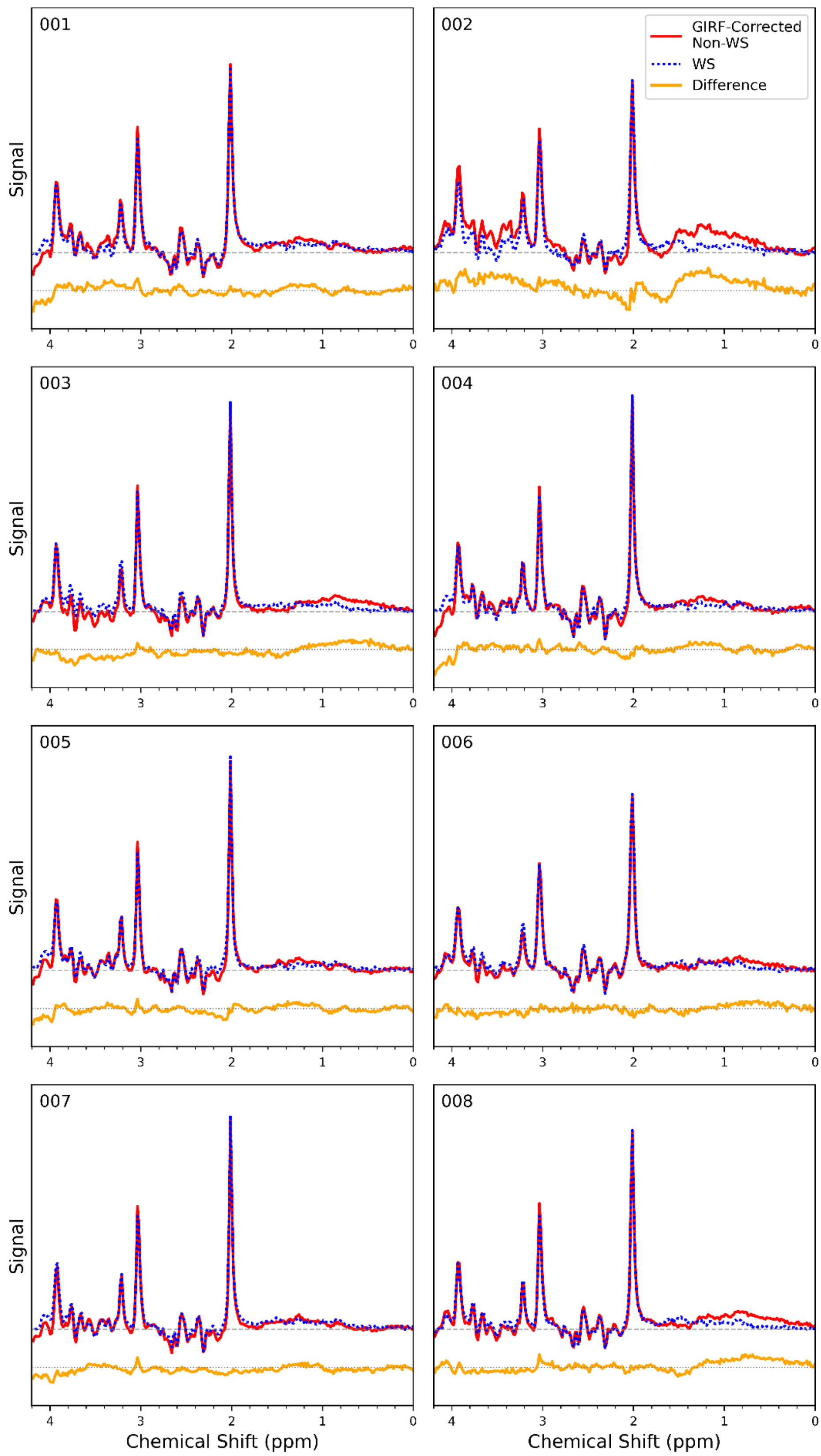


*Supporting Information Figure S1: Unfitted spectra for all participants following water removal for A) Semi-LASER acquisitions and B) MEGA-PRESS acquisitions.*

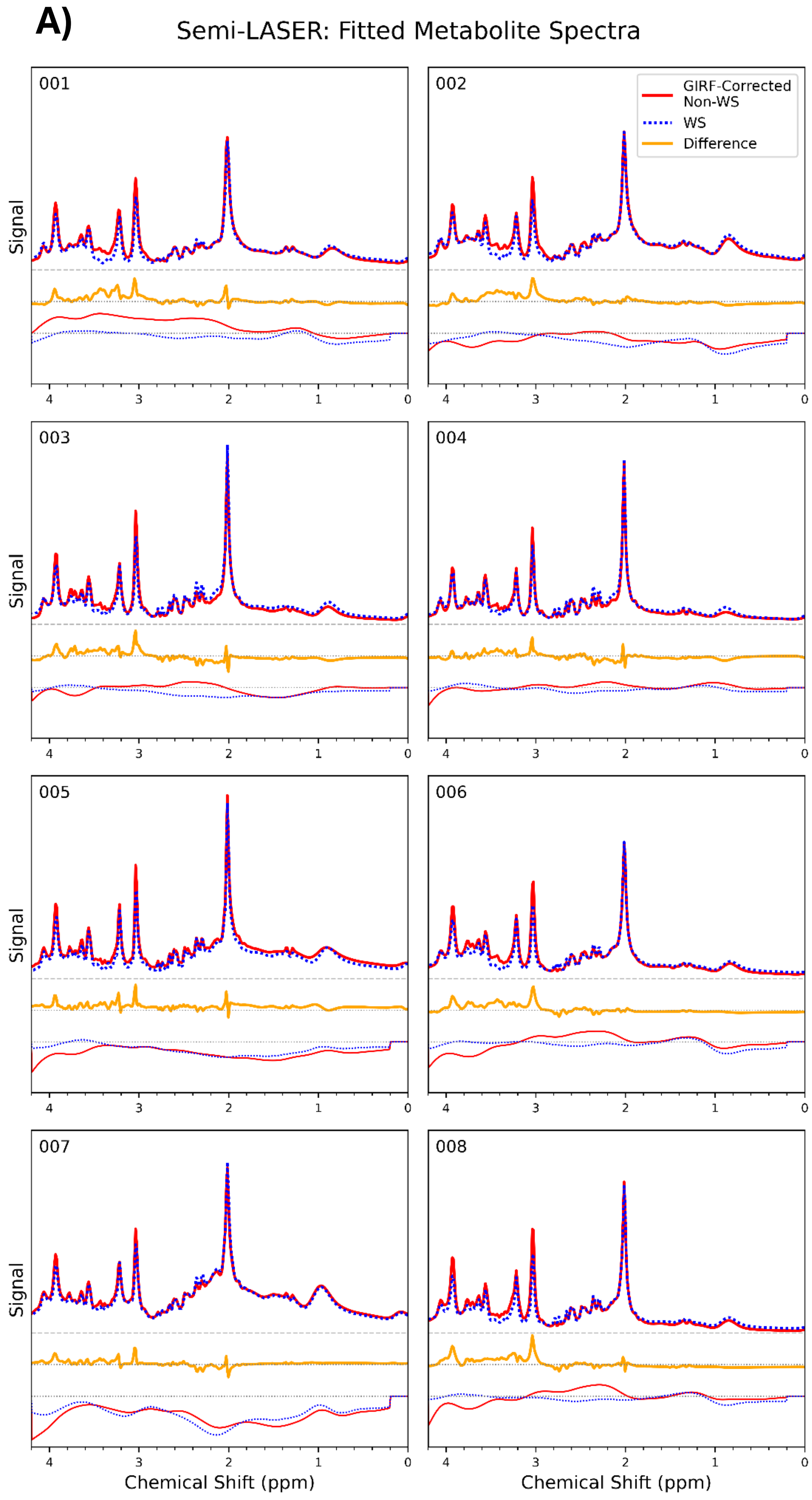
A)
Semi-LASER: Fitted Metabolite Spectra
001
002
003
004
005
006
007
008
GIRF-Corrected Non-WS
WS
Difference
Signal
Chemical Shift (ppm)
4
3
2
1
0

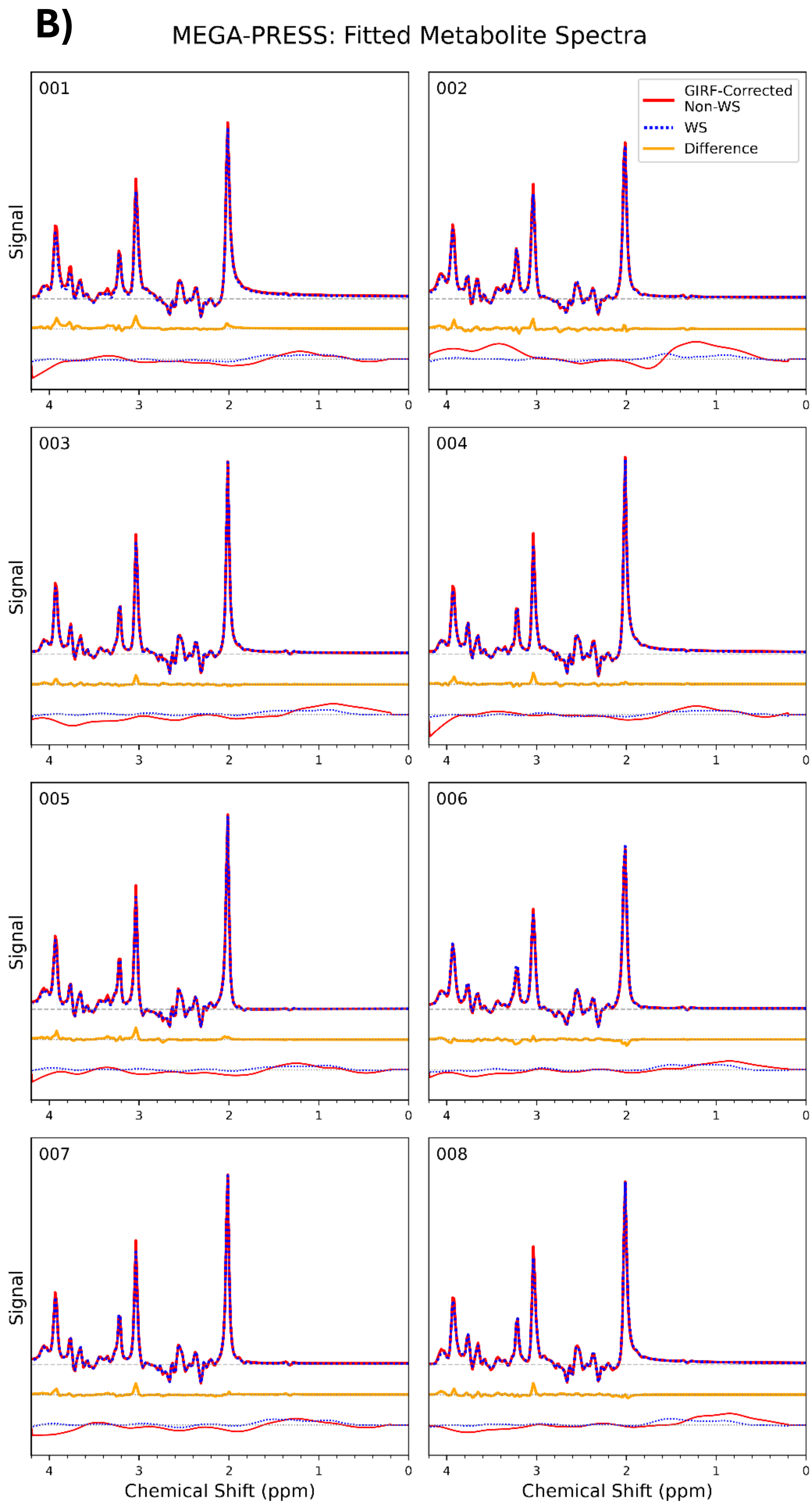


*Supporting Information Figure S2: Fitted spectra for all participants for A) Semi-LASER acquisitions and B) MEGA-PRESS acquisitions. The fitted spectral baselines are also shown and have been vertically offset for visual clarity.*

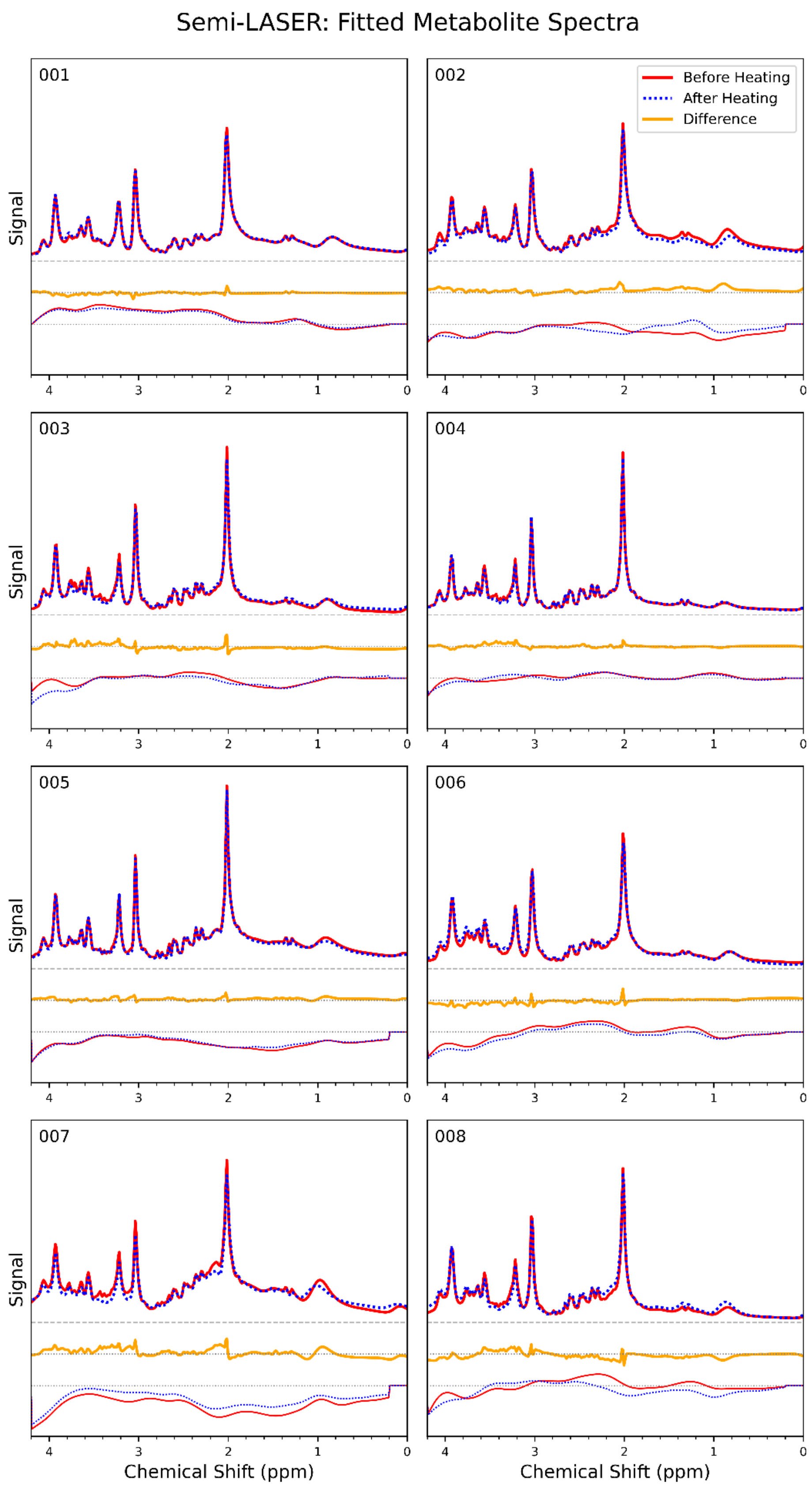


*Supporting Information Figure S3: Fitted spectra for all participants before and after system heating in the semi-LASER acquisitions. The fitted spectral baselines are also shown and have been vertically offset for visual clarity.*
